# The influence of biotic and abiotic drivers on arthropod co-occurrence network topology in native forest remnants in the Azores

**DOI:** 10.1101/2022.04.11.487852

**Authors:** Gabor Pozsgai, Pedro Cardoso, François Rigal, Mário Boieiro, Rosalina Gabriel, Eduardo Brito de Azevedo, Paulo A. V. Borges

**Affiliations:** Ce3C - Centre for Ecology, Evolution and Environmental Changes, Azorean Biodiversity Group, CHANGE – Global Change and Sustainability Institute, University of the Azores, Faculty of Agricultural Sciences and Environment, Rua Capitão João D’ Ávila, São Pedro, 9700-042, Angra do Heroísmo, Terceira, Açores, Portugal; LIBRe – Laboratory for Integrative Biodiversity Research, Finnish Museum of Natural History, University of Helsinki, P.O.Box 17 (Pohjoinen Rautatiekatu 13), 00014 Helsinki, Finland; CNRS - Université de Pau et des Pays de l’Adour - E2S UPPA, Institut Des Sciences Analytiques et de Physico Chimie pour L’environnement et les Materiaux UMR5254, 64000, Pau, France; Group of Climate, Meteorology and Global Change from the Institute of Research and Technologies for Agriculture and Environment (IITAA) from the University of the Azores. Faculty of Agriculture and Environment

**Keywords:** exotic species, network complexity, modularity, island introductions, native fauna

## Abstract

Island biota are in imminent threat from anthropogenic impacts. Of these impacts the negative effects of exotic species on the taxonomic and functional diversity of the local fauna are of particularly major concern. Aside from their impact on the diversity of native fauna, exotics may also have a detrimental effect on native interactions which, in turn, can destabilise ecological networks. Species co-occurrence networks are used to predict ecological interaction networks and utilised as tools to assess environmental impacts on community structure. Here, we investigate the topological differences of the arthropod co-occurrence networks among native forest fragments from seven Azorean islands and reveal the influence of the abiotic environment and exotic species on these networks. We found that co-occurrence networks were sensitive to environmental and community dissimilarities, showing a clear separation between islands and pinpointed differences between indigenous and exotic networks. Most exotics are little connected and exotic networks have a large proportion of unconnected species. The resulting decreased connectance and the increased modularity with the increase of the proportions of exotics in the networks suggests that most exotics have too low prevalence to show associations with other species, and only a few dominants drive co-occurrences. Moreover, the proportion of negative links, as indicators of competition, did not increase with the increase of exotics in the habitats, suggesting that exotics occupied empty niches when they colonised native forest remnants. However, when the theoretical networks consisting of only indigenous species were investigated both the number of negative associations and modularity increased with the increase of exotics, suggesting obscure competition and processes of network degradation. Since our study provides ample evidence for the usefulness of co-occurrence network analysis in studying island ecosystems, we recommend the use of this tool for ecosystem assessments, early warning systems and decision making in island biodiversity conservation.

**Significance statement:** Global anthropogenic biodiversity decline affects islands to a disproportionately greater extent than other ecosystems. One major cause of declining island biodiversity is the spread of exotic species which may overcompete and replace native biota. In this study, we show, by using arthropod species co-occurrence networks from the Azorean archipelago, that species association patterns reflect both abiotic and biotic impacts and that the increasing proportion of exotics in an ecosystem seemingly has little impact on association networks at large. However, when the effects on the association network of solely indigenous species were scrutinised, signs of network degradation were observed, suggesting an obscure, and most likely slow, negative impact of exotics on native arthropod assemblages. This disintegration of the co-occurrence networks can be the first sign of disappearing interaction links which, in turn, may jeopardise ecosystem function and can lead to regime shifts. In this work, we used a unique long-term dataset collected across the islands of the Azorean archipelago with standardised methodology. We built on the deep knowledge gathered over two decades on the ecology of species, as well as on the ongoing processes shaping the islands’ arthropod fauna, yet took a novel methodological approach and disentangled hidden ecological processes of great ecological and conservation concern.

## Introduction

Plant and animal biodiversity are declining worldwide due to human-induced stresses and insects, as the most diverse animal taxon, are critically impacted (Seibold et al. 2019, Wagner 2020, Cardoso et al. 2020, Hallmann et al. 2021). Current changes in insect species abundances with numerous extinctions are caused in different degree by habitat loss (including for agriculture), pollution (including pesticides), invasive species, climate change, direct exploitation and co-extinction of dependent species (Wagner 2020, Cardoso et al. 2020). Due to their isolated nature and fragile ecosystems, with a high number of endemic species, islands are particularly threatened by these anthropogenic stressors and thus their native species decline at an unprecedented pace (Gillespie and Roderick 2002, Fernández-Palacios et al. 2021). Whilst worldwide species declines can stem from a broad range of causes, the majority of threats to native flora and fauna on islands originates from two major sources: disappearing natural habitats due to changes in land use and the introduction of exotic species (Cardoso et al. 2010, Triantis et al. 2010, Borges et al. 2019, Pyšek et al. 2020, Fernández-Palacios et al. 2021). Whereas habitat destruction most commonly results in direct loss of species, and is relatively easy to quantify and test its effects, the consequences of spreading exotics are more difficult to study as it requires data on species interactions (Sax et al. 2002, Borges et al. 2020). These processes are thus more complex to measure, and the subtle changes can often only be unveiled by detailed community analysis. One of the first signs of such community changes is the altered network structure of species associations and interactions (Delmas et al. 2019). Indeed, interactions between species tend to break up on environmental stress sooner than species get extinct or communities change substantially (Valiente-Banuet et al. 2015). Conventional species richness or diversity-based studies may therefore be less effective in detecting changes than those scrutinizing interspecific relationships, such as associations between species pairs (Kay et al. 2018).

Undeniably, species do not live in isolation, they form ecological associations and from these associations ecological interactions emerge. These interactions underpin ecological functions, most of which are crucial in delivering the ecosystem services humans vitally depend on (e.g. Albrecht et al. 2014, Hines et al. 2015). Since biodiversity decline and homogenisation unfolds in degrading interaction networks (Laliberté and Tylianakis 2010, Burkle et al. 2013), which, in turn, decreases the stability and resilience of ecosystems and results in loss of biodiversity function (Valiente-Banuet et al. 2015), the importance of the protection of healthy ecological networks has been increasingly recognised (Tylianakis et al. 2010, Heleno et al. 2020). Hence, there is an urgent need to understand how anthropogenic impacts drive changes in interaction networks in order to precisely assess the effect of these altered networks on ecosystem functions. However, despite the considerable amount of research to investigate the anthropogenic impact on island biodiversity (Fernández-Palacios et al. 2021), little is known how anthropogenic influence impacts the interactions of species within insular communities. Different examples on how introduced exotic species can encroach indigenous network and sometimes even replace native species have been documented (García et al. 2014) though.

Since interaction networks are notoriously difficult to discover, a simple mapping of species associations based on their co-occurrence is often used as a proxy to predict interactions (e.g. Bohan et al. 2011, 2017). Although links in association networks not necessarily reflect biotic interactions (Blanchet et al. 2020), these networks nevertheless proved to be sensitive to environmental differences (Araújo et al. 2011, Lima-Mendez et al. 2015, Pozsgai et al. 2016) and to reflect anthropogenic impacts (e.g. Veech 2006, Kay et al. 2018, Elo et al. 2020). Thus, investigating relatively well-documented island faunas through co-occurrence networks offers an evident way to study how environmental factors shape local community assemblage structure and to predict the impact these factors can have on interaction networks. Analysing the structure (topology) of these co-occurrence networks can both facilitate the early detection of degrading effects and pinpointing the most vulnerable species and the most threatening exotics which, in turn, has the potential to inform stakeholders and decision-makers to maximize the success of conservation management (Delmas et al. 2019).

The Azores archipelago has been under intensive anthropogenic influence for nearly 600 years, with most of its native habitat areas being converted to agricultural landscapes (Triantis et al. 2010) and a high number of exotic species introduced (Borges et al. 2010). Taxonomic, functional and phylogenetic diversity patterns and community structures of Azorean arthropods have been widely studied (Borges et al. 2005, 2016, Rigal et al. 2018), but little attention focused on ecological networks of interspecific associations (Rego et al. 2019, Valido and Olesen 2022). Yet, the availability of this unique dataset on the arthropods of the Azores provides an opportunity to map detailed co-occurrence networks and compare them among islands and relate them to biotic and abiotic environmental factors.

We hypothesize that although species pools among the Azorean islands are highly similar, due to the presence of single island endemics the co-occurrence networks differ between islands (**H1**). We predict that island association network structure will depends on the size of habitat remnants and their proportion in the landscape as well as on the size of the island (**P1**). Moreover, island association network topologies are also likely to be driven by abiotic factors, such as temperature, precipitation, or altitude range (**P2**).

The other important factor potentially influencing the topology of co-occurrence networks is the number and the proportion of non-native species in the community. Exotics in the Azores spread rapidly (Borges et al. 2020) and although they have the potential to decrease the functional diversity of assemblages (Boyer and Jetz 2014), exotics were found to increase the functional space of native arthropod fauna in the Azorean ecosystems (Whittaker et al. 2014). The role they play in ecological networks, and how their ratio compared to native fauna influences network structure is yet to be determined. Thus, we hypothesized that association assembly between exotics and natives follows non-random organisation rules (**H2**) and that exotic species influence the structure of co-occurrence networks (**H3**). We predict that native and exotic species will not have the same role in the network, thus their node properties will differ, (**P3**) and that the topology of the theoretical networks consisting of exotic species only will have structural peculiarities (e.g. differing degree distribution) (**P4**).

Biotic stress (i.e. increased competition) can cause species to become rare, which results in these species sharing fewer sites with others and thus having fewer co-occurrence links. Ultimately, these species will not reach the detection threshold in samples and will be exempted from the networks (Kay et al. 2018). This leads to a significant decline in the number of nodes but not so much in the number of edges because of the low number of links to other species of the exempted species. Furthermore, the introduction of exotics increases the number of species (nodes) but, since they are most commonly habitat generalists, they are likely to co-occur with many other species, thus increasing the number of interspecific associations (edges) at a greater pace than that of nodes (Fridley et al. 2007). Since both processes increase the realised associations to all potential associations ratio (connectance), we predicted that the connectance will increase, and the modularity decline, with the increasing number of exotics (**P5**). We also anticipated that, if competitive exclusion is a major factor driving associations, the proportion of negative edges among all edges will increase with the increasing number of exotics (**P6**).

## Materials and Methods

Arthropod sampling followed the ‘Biodiversity of Arthropods from the Laurisilva of the Azores’ (BALA) protocol (Borges et al. 2005, 2006, 2016, Gaspar et al. 2008). Arthropods were collected in native Laurisilva forest remnants on seven islands of the Azores archipelago (Faial, Flores, Pico, Santa Maria, São Jorge, São Miguel, Terceira, Figure 1) from 1999 to 2002 (BALA I) (Borges et al. 2005) and in 2010 and 2011 (BALA II) (Borges et al. 2016). Altogether unique 91 transects were sampled, at 116 sampling occasions, 81 in BALA I and 35 in BALA II. Twenty-five sites from BALA I were repeatedly sampled in BALA II. In order to maximise the coverage of sampled diversity, two complementary methods were applied: pitfall trapping was used to sample ground-dwelling arthropods and vegetation beating was used to collect canopy-dwelling arthropods. In each forest patch, 30 pitfall traps were placed along a 150-meter long transect. Of the 30 traps, 15 were filled with Turquin (a mixture of dark beer, chloral hydrate, formalin and glacial acetic acid) and the other 15 with ethylene-glycol. In each transect, ten beating samples were taken from the three most common native woody plant species. The most common trees and shrubs sampled were *Juniperus brevifolia* (Cupressaceae), *Erica azorica* (Ericaceae), *Ilex azorica* (Aquifoliaceae), *Laurus azorica* (Lauraceae) and *Vaccinium cylindraceum* (Ericaceae) (Ribeiro et al. 2005, Gaspar et al. 2008).

**Figure 1.**
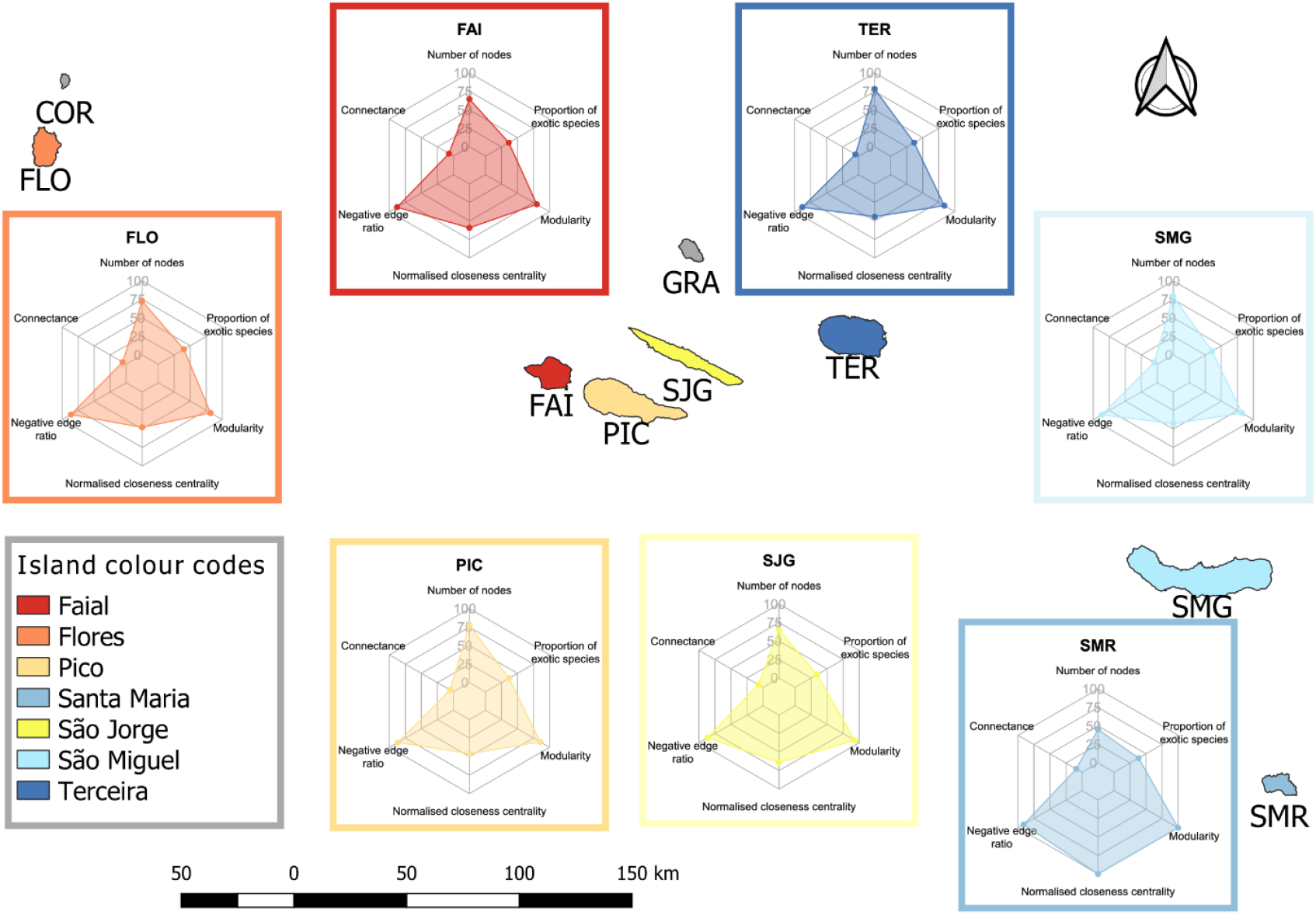
Map of the Azorean archipelago and the basic characteristics of the island co-occurrence networks. Islands are colour coded and squares with borders of corresponding colours contain radar charts showing (clockwise from the top) 1) the percentage of island species richness to the species richness in the meta-network, 2) the percentage of exotic species to the island species richness, 3) the island modularity as a percentage of the maximum modularity of all islands, .4) the normalised closeness centrality of the island as a percentage of the maximum normalised closeness centrality of all islands, 5) the percentage of negative edges of all edges in an island, 6) connectance of a network. Island abbreviations: FAI – Faial, FLO – Flores, PIC – Pico, SJG – São Jorge, SMG – São Miguel, SMR – Santa Maria, TER – Terceira.

### Environmental variables

The environmental variables were obtained from the CIELO model (Azevedo et al. 1999). This is a physical model based on the transformations experienced by an air mass crossing over a mountain, and simulates the evolution of an air parcel’s physical properties starting from the sea level up to the mountain. The model has been developed in order to produce high-resolution fields of the elemental climatic variables (pressure, temperature, rainfall, relative humidity, etc.) using a grid resolution of 100 by 100 m cell size (for more detail see also Borges et al., 2006).

### Network estimation and statistical analysis

A species-site matrix was used as a baseline dataset to generate a large meta-network, containing all species abundance data collected from all islands and sampling sites with the two sampling periods separated. When the ecological functions of adults and juvenile stages were substantially different, adults and juveniles of the same species were recorded separately (e.g. Lepidoptera). Based on Borges et al (2016), each species was categorised as either endemic, non-endemic but native (termed as native henceforth), or exotic species, and higher taxonomic levels, such as family, order, and class, were also assigned to them. Natives and endemics were sometimes merged and referred together as ‘indigenous’ species. Species whose nativity status was unknown remained included in the overall species numbers but they were not categorised into either indigenous or exotics and thus they did not inflate the number of any of those groups. Species with less than ten individuals in overall abundance and those occurring at less than three sampling sites were removed and excluded from the further analysis.

Due to methodological constrains, abundances were converted to binary (i.e. presence-absence) data and were used in the co-occurrence analysis. Two co-occurrence networks, one for the years of BALA I and one for those of BALA II, were generated using the cooccur package (Griffith et al. 2016) in an R programming environment (R Core Team 2012). The union of the two networks (i.e. merging all nodes and edges from both networks), a meta-network, consisting of species (represented as nodes) and their predicted associations (represented as links or edges) served as a base of our further analysis.

The association detecting method provided by the R package uses a probabilistic model based on hypergeometric distribution to assess if species co-occur more or less frequently than expected by mathematical chance. If a species pair occurs more often than random choice would predict, the association between them is considered as positive. On contrary, if the observed co-occurrence frequencies are lower than expected from random associations, they are considered as negatively associated. A network of positive and negative links between species nodes is thus formed by iterating through all potential species combinations. The probability for the species pair occurring together, as well as the p-values for the association being negative or positive, are given. Species and corresponding species pairs, which were expected to have less than one co-occurrence were removed prior to analysis. Species pairs with completely random association (i.e. p-value > 0.05) and those with less than 25% probability of co-occurring were removed from the generated network.

A series of 999 Erdős-Rényi random graphs with the same node number as our meta-network were generated, their degree distributions were calculated and compared to the empirical degree distribution of our meta-network to estimate the probability that our co-occurrence network is a random graph.

The meta-network was first split into seven subnetworks (henceforth island subnetworks). For each island, only species occurring on that particular island were selected to the subnetwork as nodes, and edges linking those species were retained from the meta-network. Subnetworks for each sampling site (termed as site subnetworks hereafter) were also generated, using the same method. These sub-networks were used when island networks, indigenous and exotic networks, and the node properties of native and exotic species were compared (see below).

Commonly used measures to characterise network topological properties, such as the number of nodes and edges, connectance, proportion of negative links, the proportion of isolated nodes, mean closeness and betweenness centralities, and modularity (based on the ‘fast and greedy’ community detection algorithm) were calculated. Similarly, node characteristics, such as the number of edges connecting other nodes (degree), the proportion of degree to the number of all nodes (relative degree); the number of negative edges (vulnerability) and the proportion of those to the relative degree (relative vulnerability); betweenness, closeness centralities were also computed, with the help of the igraph package (Csárdi and Nepusz 2006). Since most of these measures strongly depend on the number of nodes (i.e. show high correlation), a z-score normalised version of each centrality measure was also calculated. During the process of excluding highly correlating variables (Spearman’s p<0.05, Spearman’s Rho> 0.6 or Spearman’s Rho<-0.6) non-normalised versions of these variables were discarded and only normalised values were included in the analysis. The process of network generation and the description of all calculated network measures, the way how they were calculated, and their correlations are given in Supplementary Material 1.

All network properties were calculated for each subnetwork, and node properties were calculated in each subnetwork separately to all species, indigenous and introduced.

Site networks were used to compare island networks and to investigate their relationship to the island area, the total native forest area, and the native forest to island area ratio, as well as to island-specific climatic variables and the number of exotics and their ratio to the total species richness on the islands. Significantly highly correlating environmental variables (Spearman’s p < 0.05, Spearman’s Rho > |0.6|) were removed prior to the analysis. Since there is no settled methodology to compare networks to each other, two different approaches were taken: 1) island networks were compared based on their associated species pairs, where distance matrices were calculated using the Jaccard distance on presence-absence matrices of species associations; and 2) based on the differences in their calculated network properties. In this latter case, similarity matrices were calculated using the Euclidean distances of z-score scaled network properties. Environmental variables were also z-score scaled and a stepwise redundancy analysis (dbRDA) process was conducted to find the optimal model. Whether or not island networks topologies were significantly different was tested using the corresponding distance matrices in an Analysis Of Similarities (ANOSIM) test with 10000 permutations, with the help of the *anosim*() function implemented in the vegan (Oksanen et al. 2010) R package. To controlling Type I Error arising from multiple comparisons, p-values in pairwise comparisons were adjusted using the false discovery rate (FDR) correction.

To test if link formation between either combination of endemic, native, and introduced species occurred non-randomly, we generated 5000 networks in an iterative process with keeping the original network structure but randomly assigning the origin status to species. The proportion of each combination pair to the overall link number was calculated and one-sample t-tests were used to compute p-values to determine if association frequencies between categories of native status can be random. A similar permutational approach was used to test whether or not some combinations of categories of native status are more or less likely to collect negative links than it would be expected from random processes. In this latter case though the ratio of the negative links to the number of links within each combination pair was calculated and compared to the randomised distribution, using one-sample t-tests. To investigate if there are differences in the frequency of endemic, native, and introduced species having association links with each other we compared the number of links between each combination using Kruskal-Wallis tests, and pairwise Wilcox-tests with p-values adjusted according to the FDR method.

The relationship between major network topology measures and the number and the proportion of exotics in the communities was investigated using linear mixed models with the island identity set as the random term using the lmer() function in the lme4 package (Bates et al. 2015 p. 4). All proportion variables were square-root transformed prior to regression to approximate normality. P-values were estimated according to Satterthwaite’s method, as implemented in the lmerTest package (Kuznetsova et al. 2017) in R. Marginal and the conditional R2 values were extracted using the *r*.*squaredGLMM*() function in the MuMIn package (Barton 2020). Since indigenous and exotic species richness strongly correlated, to better disentangle the effects of exotics, similar models as above were fitted on networks consisting of native species only and residuals from these models were used to re-fit models between node properties and the number and proportion of exotics.

Both network-related and node-related properties were compared between native and exotic species using Kruskal-Wallis tests and pairwise Wilcox tests with p-values adjusted for multiple testing according to the FDR method. Linear mixed-models as above were used to investigate the effect of exotics (both number and proportion in the whole community) on indigenous networks only.

## Results

Our initial meta-network consisted of 161 nodes (species) and 398 edges, giving a 0.031 edge density value. Of all species, 101 (52 endemics and 49 natives) were indigenous, 58 exotic, and 2 with an unknown origin. Positive associations overwhelmingly dominated the meta-network (345 and 53 positive and negative links, respectively). The edge density and degree distribution of our meta-network were significantly different from those that could have arisen from random networks. The degree of nodes ranged from 1 to 44 with *Argyresthia atlanticella* Rebel, 1940, a moth species endemic to the Azores, having the highest degree. *Lasius grandis* Forel, 1909, a native ant, and *Palliduphantes schmitzi* (Kulczynski, 1899), an endemic spider, had the most negative link to other species (Figure 2, Supplementary material 2).

**Figure 2.**
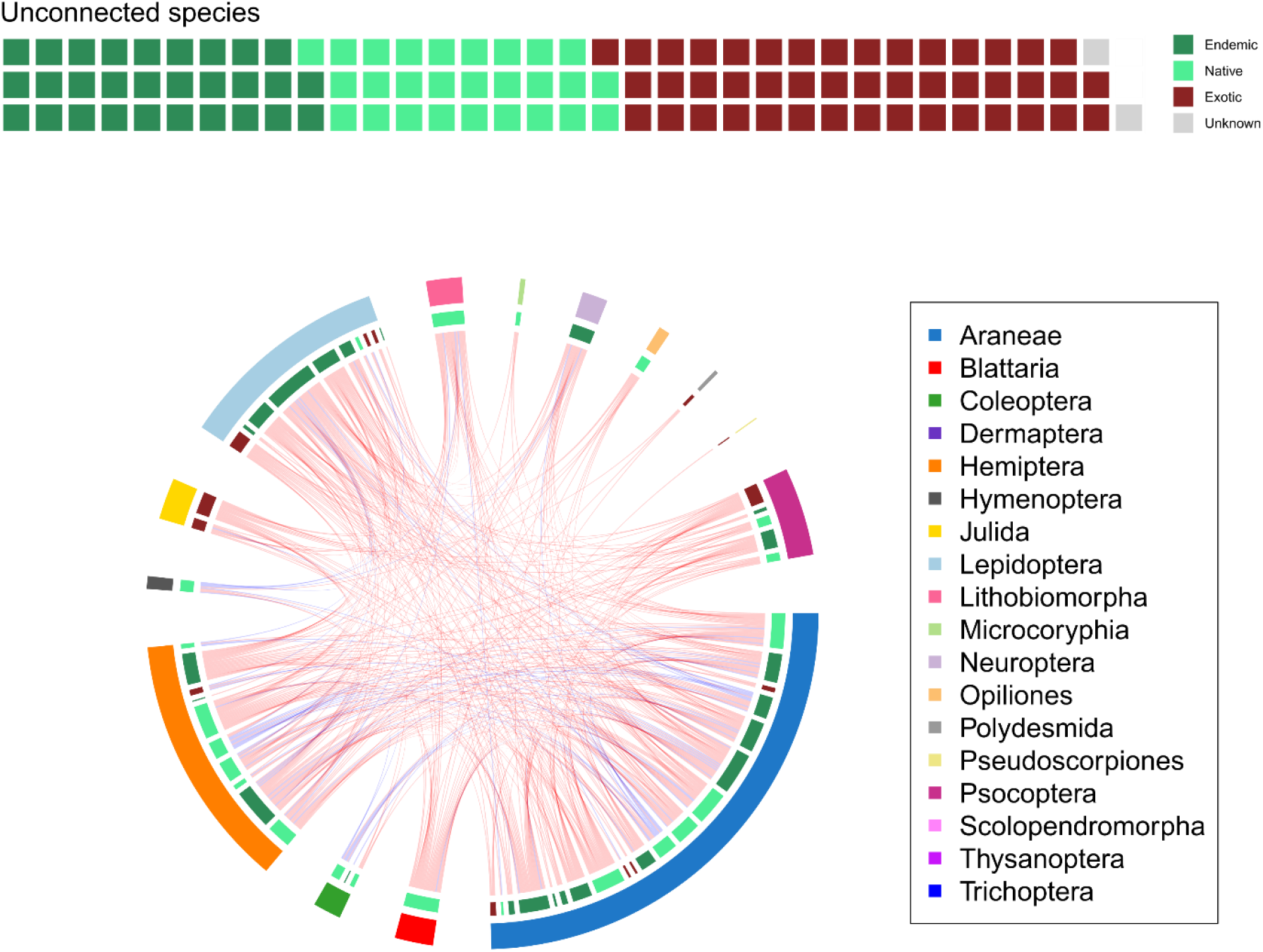
Meta-network coloured according to higher taxa and nativity classes (A), and the number of isolated species, grouped to endemics, natives, exotics, and unknown origins (B). Red links in the network represent positive, blue links negative associations. Outer arc shows arthropod orders, inner arc nativity classes. Arc segment length is proportional to the number of nodes the group has.

### Differences in island network topologies and driving factors

Islands networks significantly differed based on their topology measures (ANOSIM p <0.001, R = 0.175). After correcting for multiple comparisons, several pairwise differences between islands still remained significant at the p < 0.05 significance threshold (Figure 3A-B). Although the number of nodes and the number of edges were significantly different between islands (Kruskal-Wallis test, p = 0.04 and p = 0.02, respectively), pairwise differences were not supported statistically. The connectance, the ratio of the isolated nodes to all nodes, the relative vulnerability, normalised closeness and betweenness centralities, and modularity, on the other hand, showed significant pairwise differences (Figure 1B, Supplementary material 3). The number and proportion of the exotics on the islands, modelled mean altitude, and the annual mean precipitation were the main factors driving these differences (dbRDA model was significant at the p=0.003 level and explained 15.9% of constrained inertia) (Figure 3C). When island networks were compared based on their association pairs, they differed significantly (ANOSIM p <0.001, R = 0.229) but differences between individual islands were different than those when island networks were compared based on their topology measures (Figure 3D). The major factors driving these differences were the area of the native forest on an island, annual mean, and summer median temperatures, and summer precipitation and average relative humidity (dbRDA model was significant at the p<0.001 level and explained 19.7% of constrained inertia).

**Figure 3.**
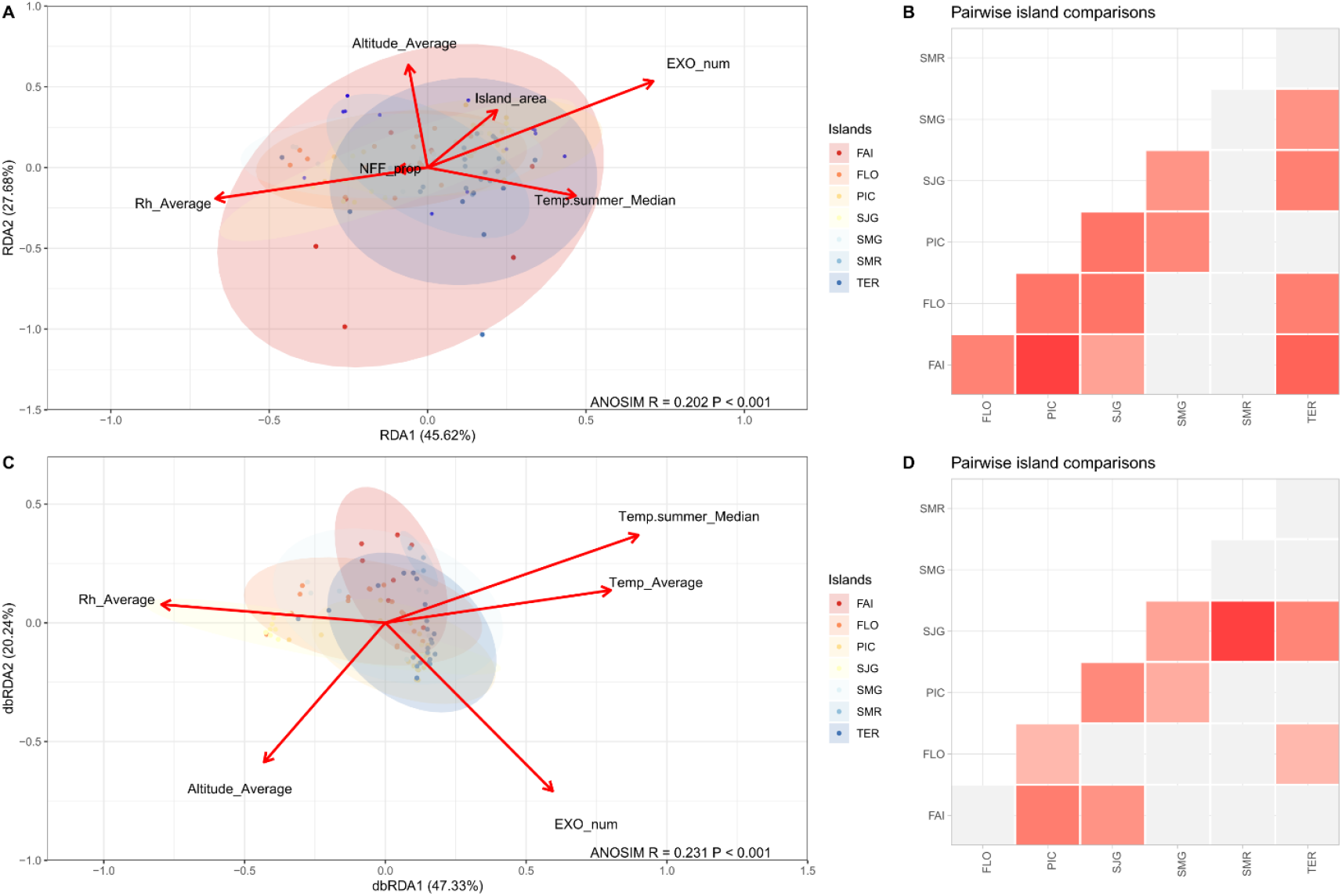
The ordination and pairwise ANOSIM comparison of the island subnetworks based on their network properties (A, B, respectively) and their species pair community (C< D, respectively).

### Effects of exotics on network topology

Simulations suggested that endemic to endemic, endemic to native, and native to native edges were less common in the meta-network than could have arisen in networks with randomly reshuffled nativity categories. At the same time, introduced species were more linked to the other groups than expected by chance, including themselves. Moreover, natives had a lower chance than expected to have negative links to both endemics and other natives. All other combination pairs showed a significantly greater chance than random to have a higher proportion of negative links with each other (Supplementary material 4). When linking frequencies were compared, endemic to endemic, native to native and endemic to native links occurred in greater proportions than exotic to exotic, and exotic to native/endemic (Figure 4A). In terms of the proportion of negative links, natives had negative associations with themselves or with the other two categories in significantly higher proportions than any other combinations. Exotic to exotic negative links occurred in a significantly lower proportion than endemic to endemic ones (Figure 4B).

**Figure 4.**
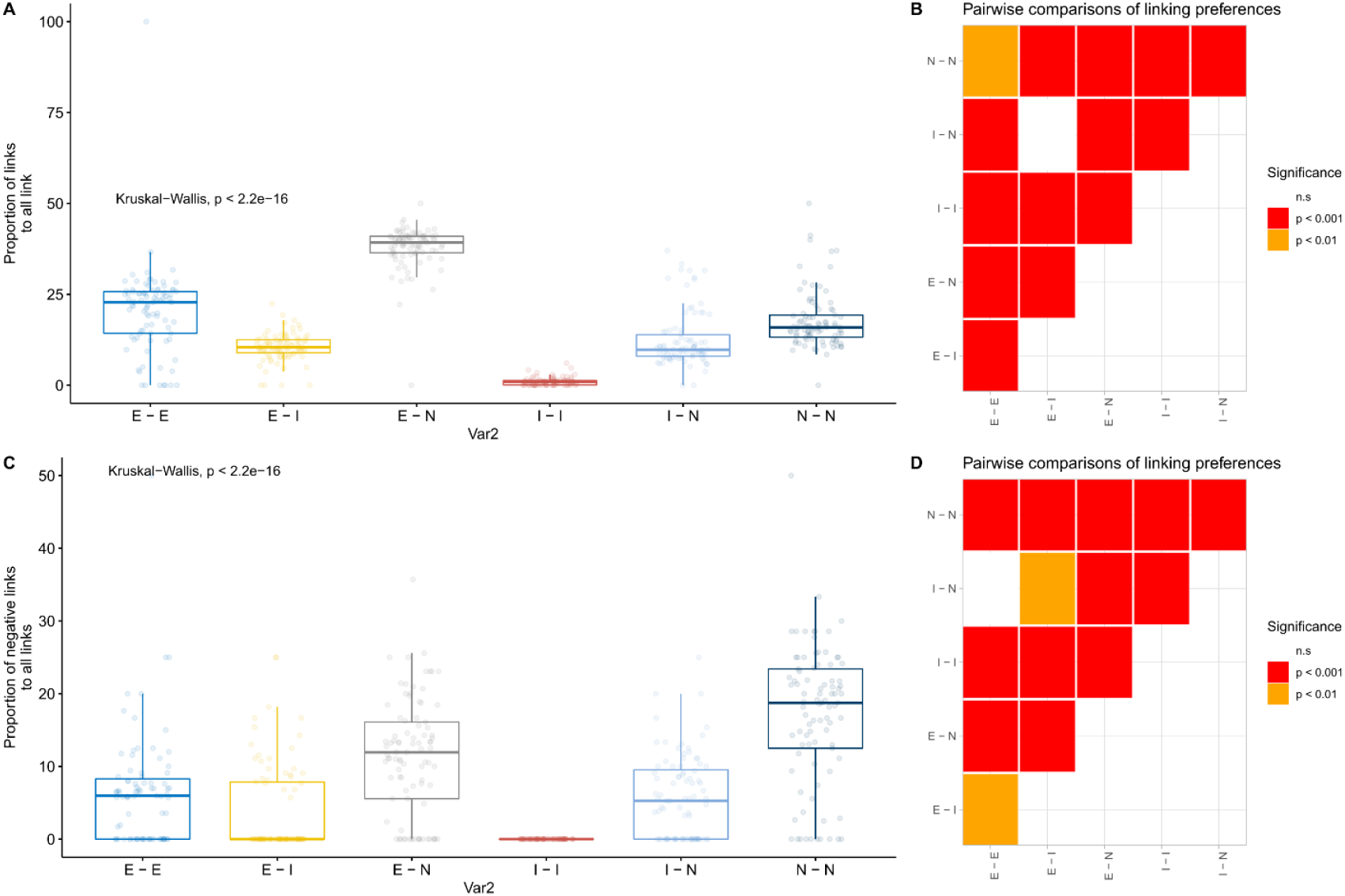
Preferential linking between endemic, native, and exotic species. The distribution of the proportion of both positive and negative links (A), and negative links only (C) to all links between nativity category pairs. Pairwise comparisons using pairwise Wilcox tests are shown on the right side (B, D, respectively). All p-values of pairwise Wilcox tests are adjusted according to the FDR method.

Both the number of nodes and edges, as well as the ratio of negative edges showed significantly positive relationships with the number of exotics in the networks. On the contrary, the connectance, and the normalised closeness and betweenness centralities showed negative relationships (Table 1, Supplementary material 5).

**Table 1.** R^2^, f, and p-values for each network parameter as a function of the number and proportion of exotics in the community. Models ran for all site subnetworks, site subnetworks consisting of indigenous species only, and for the residuals of the model fit on indigenous species against the number of exotics.

|  |  | Number of<br>nodes | Number of<br>edges | Proportion<br>of isolated<br>nodes | Proportion<br>of<br>negative<br>edges | Mean<br>degree | Connectan<br>ce | Normalise<br>d<br>closeness<br>centrality | Normalise<br>d<br>betweenness<br>centrality | Modularit<br>y |
| --- | --- | --- | --- | --- | --- | --- | --- | --- | --- | --- |
| **Full network** | R<sup>2</sup> | 0.676 | 0.371 | 0.173 | 0.289 | 0.217 | 0.445 | 0.303 | 0.254 | 0.261 |
| **–** | F | 159.371 | 29.589 | 3.814 | 12.493 | 2.417 | 63.17 | 38.641 | 4.093 | 1.392 |
| **exotic**<br>**number** | P | 0 | 0 | 0.054 | 0.001 | 0.124 | 0 | 0 | 0.046 | 0.241 |
| **Full network** | R<sup>2</sup> | 0.232 | 0.374 | 0.696 | 0.251 | 0.649 | 0.336 | 0.206 | 0.429 | 0.376 |
| **–** | F | 14.333 | 42.992 | 149.246 | 0.029 | 136.697 | 24.694 | 8.537 | 25.487 | 19.625 |
| **exotic**<br>**proportion** | P | 0 | 0 | 0 | 0.866 | 0 | 0 | 0.004 | 0 | 0 |
| **Native** | R<sup>2</sup> | 0.36 | 0.291 | 0.35 | 0.256 | 0.227 | 0.392 | 0.307 | 0.37 | 0.206 |
| **network –** | F | 41.103 | 15.74 | 5.89 | 0.058 | 2.807 | 45.073 | 30.493 | 4.241 | 4.446 |
| **exotic**<br>**number** | P | 0 | 0 | 0.018 | 0.81 | 0.098 | 0 | 0 | 0.043 | 0.038 |
| **Native** | R<sup>2</sup> | 0.282 | 0.38 | 0.455 | 0.378 | 0.521 | 0.254 | 0.179 | 0.308 | 0.275 |
| **network –** | F | 26.842 | 42.484 | 26.001 | 20.954 | 70.384 | 8.471 | 8.905 | 1.123 | 19.351 |
| **exotic**<br>**proportion** | P | 0 | 0 | 0 | 0 | 0 | 0.005 | 0.004 | 0.293 | 0 |
| **Full network** | R<sup>2</sup> | 0.597 | 0.063 | 0.199 | 0.034 | 0.233 | 0.197 | 0.106 | 0.065 | 0.128 |
| **residuals –** | F | 117.742 | 5.992 | 22.078 | 3.112 | 27.022 | 21.813 | 10.502 | 6.162 | 13.088 |
| **exotic**<br>**number** | P | 0 | 0.016 | 0 | 0.081 | 0 | 0 | 0.002 | 0.015 | 0 |

However, when relationships between these measures and the proportion of exotic species in the community were tested, only the proportion of isolated nodes and modularity revealed significantly positive relationships but the number of nodes and edges, the mean degree, the ratio of negative edges, the normalised betweenness centrality, and the connectance showed significantly negative relationships. Relationships showed similar patterns when the proportions of exotics were fitted against the residuals of the model fitted on the proportions of exotics against the properties of indigenous-only networks (Table 1, Supplementary material 6).

### Differences between the node properties of indigenous and exotics

The mean and the relative degree, the normalised closeness and betweenness centralities, as well as the number and ratio of positive links to other species, were greater for indigenous species in the meta-network. However, indigenous and endemics only showed significant differences in their normalised closeness centrality once isolated nodes (degree=0) were removed (Supplementary material 7)

### Differences in native and exotic networks

When island subnetworks were split into networks consisting of only indigenous or exotic species, differences emerged. Since there were more indigenous species than exotic, network topology measures highly correlated to node number (such as number of edges, number of positive links, mean degree etc.) were also significantly greater for native networks. Albeit they had no or little correlation to the number of nodes, the proportion of isolated nodes, the connectance, and the normalised closeness and betweenness centralities also showed differences (Figure 5).

**Figure 5.**
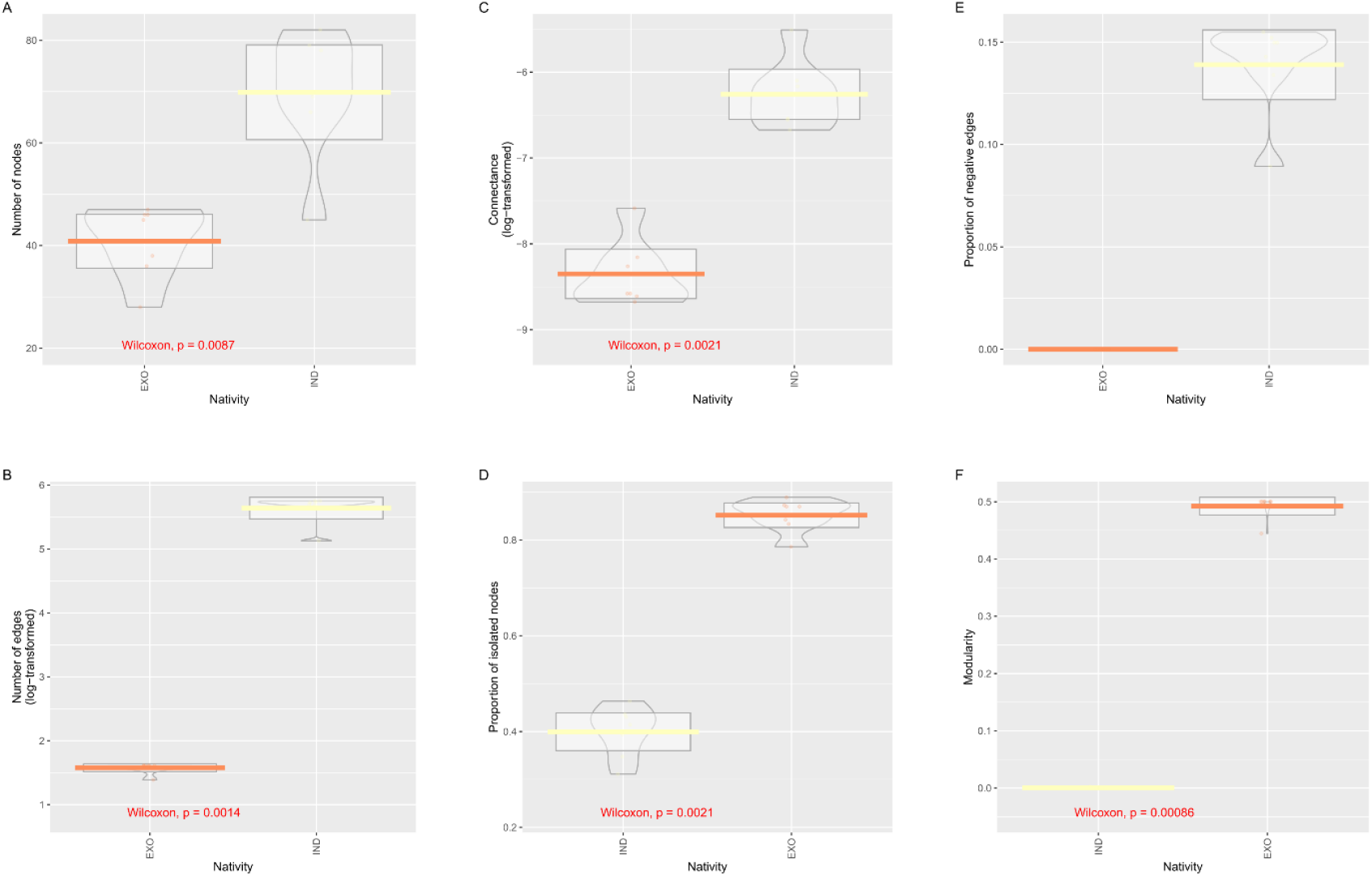
Comparison of network properties between indigenous-only and exotic-only networks: number of nodes in the network (A), number of edges in the networks (B), the log-transformed value of connectance (C), the proportion of the isolated nodes (D), the proportion of negative edges (E), and the modularity based on the ‘fast and greedy’ community detection algorithm (F). Note that the subnetworks in the figures are ordered according to the mean value of the focal network measure.

When we investigated the effects of exotics on indigenous networks only, we found that number of nodes (species number) and edges decreased with increasing exotic proportion. Exotics had a similarly negative relationship with the mean degree and the normalised betweenness centrality. A positive relationship was visible between the proportion of exotics and the proportion of isolated nodes, the connectance, the normalised closeness centrality, and the modularity (Supplementary material 8).

## Discussion

In this study, we analysed arthropod co-occurrence networks on seven Azorean islands and tested the hypotheses that these networks reflect biogeographical patterns, are sensitive to abiotic environmental differences, and that their topological features echo the imprint of exotic species in the community.

We found that co-occurrence networks of island arthropods showed non-random structuring patterns, and that biogeography (i.e. island identity) was reflected on the network structure both when species pairs, as network building blocks, and when network topological properties were compared (H1). Both of our first and second predictions (P1, P2) that natural habitat size and abiotic factors drive network structure in concert, were supported by the multivariate model. Thus, co-occurrence network analysis seems to be suitable to detect inter-island differences and the dependence of network topology on environmental factors is clear. Yet, although because island species richness strongly depends on the size of the island (Whittaker et al. 2017) and natural habitat remnants behave as islands themselves (Matthews 2021), we predicted natural habitat patch size will influence the structure of the association networks, our results showed that the size of natural habitat has lower importance in shaping co-occurrence networks than they have in driving community differences in indigenous Macaronesian spiders (Cardoso et al. 2010). However, Cardoso et al. (2010) excluded exotic species from their analysis, and in our cases, the number of exotics dominated the model, thus the disagreement with their findings can be explained. Moreover, in our study, there was a moderately strong correlation between native forest patch size and the number of exotics, which may have further obscured the clear effect of forest patch size. Nevertheless, native habitat area and proportion showed a few, moderately strong, correlations with network properties (Supplementary material 9), suggesting a limited power of this variable to predict networks topologies.

Both the number and the proportion of exotics in the community influenced the structure of co-occurrence networks (H3). As suggested by the species-area relationship (Whittaker et al. 2017), the number of nodes showed a positive correlation with the island area, and so did the number of exotic nodes. This is in line with the findings of Whittaker et al. (2014), who reported an increasing number of exotics with increasing island area for both spiders and beetles in the Azores. The proportional increase of edges did not match the increase of exotic nodes though, mostly because the newly recruited exotics in the communities have no or few links to other species (i.e. the proportion of isolated increased). This resulted in a general decline in the connectance, and, when the proportion of exotics to the entire community was investigated, an increase of modularity; the opposite way we predicted (P5). These trends are more pronounced when the effect of indigenous species is removed, suggesting that indigenous mitigate changes in association network structure. Moreover, the declining number of nodes as a function of the proportion of exotics in the community suggests uneven recruitment of new species into the communities: when species richness increases, newcomers are mostly exotics. The fact that the proportion of negative links did not show a significant relationship either as a function of the residuals after the effect of natives had been removed, or when the proportion of the exotics was investigated (i.e. P6 did not hold up), suggests that these species are rarely involved in direct competition with indigenous ones. This pattern, and the high proportion of unconnected exotic species, on one hand, suggest that the majority of the exotics do not occur in samples regularly enough to form detectable associations with other species; only a few, dominant, exotics contribute to shaping network topologies (Kay et al. 2018). This is in line with Florencio et al. (2015) who found that faunal homogenisation in the Azores was not apparent from incidence-based community nestedness investigations, and reasoned that although the prevalence of dominant exotic species was high, rare exotic species were replaced both in space and time. On the other hand, our results support the earlier findings (Whittaker et al. 2014) that exotics, instead of competing with indigenous, occupied empty niches and increased the realised trait space of the community (Rigal et al. 2018). However, the increasing proportion of negative associations between indigenous species with the increasing proportion of exotics suggests an increasing indigenous to indigenous competition as the effect of exotics.

We also showed a strong preferential linking in the community, and consequently, the assembly structure was not random (H2). Endemic and native species linked to each other more frequently than to exotics. This is somewhat controversial to what we expected, that since exotics are habitat generalists and occur in many habitats, they will regularly co-occur with all species, and thus have a high number of links (including negative ones). Similarly to the previous section, the reason for this may be the relatively low number of exotic species being prevalent enough for association detection. Indeed, although native habitat fragments are relatively small, most many exotic species may not reach the locations toward the centre of patches where indigenous are frequent. Whether this happens through the resistance of local communities to exotics or other reasons is yet to be investigated. Moreover, the number of endemic to endemic links may have been inflated through species turnover within archipelago due to speciation. Preferential linking through negative links was not obvious either and the trend in the proportion of negative links in communities was also unclear (P6), suggesting little niche overlap and competition to indigenous species in the Azores (e.g. Heleno et al. 2013).

Networks consisting of solely indigenous or exotic species also differed, as we predicted (P3). Exotic species had different node properties than indigenous, but they showed a generally lower number of links to other species and the proportion of negative links showed a significant relationship with the number, but not with the proportion of exotics in the community (Thus, P7 was only partially upheld.). This low degree resulted in lower connectance and centralities, and a greater proportion of isolated nodes in exotic-only networks, compared to native-only networks (P4). As a consequence, connectance, indeed, decreased and modularity increased with the increase of the number and proportion of exotic species in communities (P5). Although, as seen above, these can be the results of exotic species blending into indigenous communities without competing with indigenous species, from the high modularity of exotic networks we also may speculate to their lower stability. Indeed, as a number of systems show early signs of disintegration when stressed, particularly the weak links tend to break easily (Csermely 2004), increasing modularity is anticipated. Alarmingly, in our native-only networks, the modularity also decreased with the increasing proportion of exotics in the community, as did the proportion of isolated nodes and negative links. These suggest an obscure process of disintegration of native association networks, driven by the increasing proportion of exotics, which, eventually may grow into a regime shift (Rocha et al. 2015, Hui and Richardson 2018). This is in line with, Larson et al. (2016) and Hui (2021) who showed that plant-pollinator interactions and fruit-bird mutualistic networks (respectively) change in a similar manner when invaded by introduced species. Although co-occurrence networks cannot be translated to interactions (Blanchet et al. 2020), species pairs that do not co-occur cannot interact either, and hence these findings are highly concerning and in accordance with the recent observation that exotic species diversity is increasing in Azorean native forests (Borges et al. 2020). Moreover, the number of nodes was declining with the increasing proportion of exotics but the connectedness increased, indicating that less connected species disappeared first, reinforcing the estimations by Triantis et al. (2010) for a high level of extinction debt on the Azores.

Nonetheless, since species occurrences may also correlate with latent environmental factors, for instance, the adjacent landscape of the natural forest patch, other drivers may also be in action. Thus, before drawing casual links between exotic species’ number and node properties and native species richness, the underlying causes should be thoroughly investigated.

Our study provided ample evidence that island arthropod co-occurrence networks are sensitive to the presence of exotic species and that the networks of exotic species differ from those of natives. These structural sensitivities can make species co-occurrence networks ideal tools for providing early warning signals of community changes induced by exotics. These signalling systems in the Anthropocene are timely and essential to detect and mitigate deleterious effects of human-induced environmental change on native habitats (Derocles et al. 2018, Fath et al. 2019). On the other hand, in the last decades, the amount of biodiversity data multiplied, partly due to the advanced recording technology (e.g. metabarcoding, environmental DNA), but also due to citizen science efforts. These untapped data could be utilised for co-occurrence network analysis to understand large-scale ecological assembly rules and geographic patterns of communities (Lima-Mendez et al. 2015, Ma et al. 2016) as well as for early warning systems in conservation. A cautious approach has to be taken though. In our case, for instance, negative links between species did not provide a useful measure for the effect of invasive species, most likely because, as we speculated, the exotic arthropods on the Azores naturalised relatively well and managed to exploit previously unoccupied niches causing little competition with natives, as it was reported in the case of disturbed landscape such as managed pastures (Rigal et al., 2018). Whether or not this process drives the patterns we found in native forests, can only be teased apart through targeted field experiments.

### Limitations

One of the main limitations of this study is inevitably derived from limitations of the method used; although association networks are relatively easy to construct, they are not real-life interaction networks, merely the predictions of them (Blanchet et al. 2020, Strydom et al. 2021). This is particularly true because co-occurrence networks are scale dependent; although our sampling transects were relatively small (150m), less mobile or microhabitat restricted species are unlikely to interact even at that spatial scale. Therefore, a deeper insight is needed into the pairwise links and targeted tests or literature searches should prove or disprove the existence of predicted interactions. Although the dynamism of these networks is accounted for in our study (two separate networks were generated for the two sampling rounds), deep dynamical processes are not analysed. This limitation is the direct consequence of the lack of underpinning long-term datasets. This deficiency restricts our understanding of processes overarching several decades, such as climate change, the temporal patterns of exotic invasions, or continuous anthropogenic pressure, and likely prevents timely action to mitigate them (Poisot et al. 2015, Tulloch et al. 2016). Moreover, species co-occurrence networks may also depend on the seasonal dynamics of species of which we have little information. In this study, we did not focus on differences resulting from taxonomical or functional grouping but these, most likely, exist. Whereas this approach would plausibly be a fruitful area of research, a complete dataset of traits is crucial and, besides taxonomy, a phylogenetic tree would also be desirable.

## Conclusions

Here we show that changes in the topologies of arthropod co-occurrence networks in the Azores mirror variances both in biotic and abiotic environments and thus they can help to gain a deeper insight into natural and anthropogenic processes shaping island biogeography. Our findings demonstrate that although Azorean exotic species have little competition to indigenous, their presence affects species association networks and induce alarming reorganisations. Thus, developing standardised network assessment methods and utilizing network information may help in developing early warning systems for detecting the perilous impact of exotic species (Fath et al. 2019). Combining modern metabarcoding techniques and standardised statistical methods for association network-building with cutting-edge machine learning processes and literature-based trait data to routinely identify real-life interaction networks would substantially advance our understanding of ecological assembly rules and improve our predicting power to anticipate the future status of communities of high conservation interest (Evans et al. 2016). Fully exploiting this toolkit is vital for island biodiversity conservation.

## Author contributions

G.P. and P.A.V.B. conceived the idea. Data were collected by the contributors of the BALA dataset (Borges et al. 2016) and the environmental data were provided by E.B.A. Data analysis was conducted by G.P., who also wrote the first version of the manuscript. The first draft was edited by P.C., F.R., M.B., and P.A.V.B. The necessary funds were acquired by P.A.V.B. All authors contributed to editing the manuscript. All authors have read the manuscript and agreed with its content.

## Data archiving statement

Species list and distribution data are openly available in Borges et al. (2016). Computer codes, along with summary environmental data will be made available on GitHub upon publication.

## Acknowledgements

We are grateful to all colleagues that contributed to data acquisition during BALA 1 and BALA 2 fieldwork.

**Supplementary material 1. Network measures calculated in the study and their ecological importance**.

**Supplementary material 2. List of all species in the meta-network with their taxonomic grouping (Order, Family), nativity categorisation, and calculated node measures**.

**Supplementary material 3. Comparison of network properties of island subnetworks. Pirate plots show the distribution of the data, the median, and the lower and higher quartiles. Kruskall-Wallis tests are displayed, and pairwise differences are also shown if Wilcox tests are significant after adjusting p-values with the FDR method**.

**Supplementary material 4. Preferential pairing between endemic, native, and exotic species. P-values of the one-sided t-tests comparing the measured and expected values from the simulation process are shown for all links and negative links only (A). The results of proportion tests between the frequencies of links between exotics – natives and natives – natives (B), exotics – endemics and endemics – endemics (C), and exotics – indigenous and indigenous – indigenous (D) species**.

**Supplementary material 5. Relationship between the number of exotic species and major network properties in networks consisting of all species**.

**Supplementary material 6. Relationship between the number of exotic species and the residuals of the model fitted against the major network properties in networks consisting of indigenous species only**.

**Supplementary material 7. Differences between the node properties of indigenous and exotic species**.

**Supplementary material 8. Relationship between the number of exotic species and major network properties in networks consisting of only native species**.

**Supplementary material 9. Correlation between all network properties, environmental variables, and between network properties and environmental variables**.

