## Supplementary Material 1 for "The influence of biotic and abiotic drivers on arthropod co-occurrence network topology in native forest remnants in the Azores"

| | | | |
|---|---|---|---|
| **Network properties** | | | |
| Number of nodes (n) | The sum of all nodes present in the network | Equals to species richness. | $N = \sum n$ |
| Number of edges (e) | The sum of all association links between species (nodes). | All associations between species pairs. | $L = \sum e$ |
| Connectance | The proportion of realised links to all possible links. | Shows how dense is the association network. Does not depend on species number. | $\frac{L}{N(N-1)/2}$ |
| Mean degree | The average number of links per node in the network. | Shows how dense is the association network. Depends on species number. | $\frac{\sum_{i=1}^{N} d_i}{N}$ |
| Negative edge ratio | The proportion of negative edges in the network to all edges in the network | Shows what proportion of between-species associations are negative, i.e. most likely competitive | $\frac{\sum_{i=1}^{L} e_{i,sign=-1}}{L}$ |
| Proportion of isolated nodes | The proportion of nodes with no links to other nodes to the number of all nodes in the network | Shows what proportion of species in the community does not have association links with other species. | $\frac{\sum_{i=1}^{N} n_{i,d=0}}{N}$ |
| Modularity | Sum of the differences between the between-module edges (*e<sub>ij</sub>*) and the fraction of within-module edges (*a<sub>i</sub>*) in the community. | Indicates to what extent the network is divided to groups of species that have more within-group links thank links to outside of the group (modules). | $\sum_{i=1}^{c}(e_{ij-a_i^2})$ |
| **Node properties** | | | |
| Degree | The number of links a node ($n_i$) has to other nodes | Shows how many other species one species has association links. Likely to correlate with the number of interactions. | $d_i = \sum e_i$ |
| Relative degree | The proportion of the degree of the node to the number of all edges in the network | A measure of how well connected a species is in the association network. Independent of species richness (number of nodes in the network) | $\frac{d_i}{L}$ |
| Vulnerability | The number of negative links a node has to other nodes | Number of species with which the focal species has negative associations, most likely competitive interaction. | $V_i = \sum_{i=1}^{d_i} e_{i,sign=-1}$ |
| Relative vulnerability | The proportion of negative links the node has divided by the node's degree | A measure of the importance of negative associations for a species. | $\frac{V_i}{d_i}$ |

| | | | |
|---|---|---|---|
| Normalised betweenness centrality | The number of shortest paths between two nodes passing through. | Quantifies the functional importance of a species affecting other species. | $C_C(P_i) = \frac{N-1}{\sum_{k-1}^{N} d(P_i, P_k)}$ |
| Normalised closeness centrality | Sum of distances from other nodes. | Quantifies the functional importance of a species as a connector between parts of the network. | $C_B(n_i) = \sum_{j<k} g_{jk}(n_i)/g_{jk}$ |
