## Supplementary figures and images for "The influence of biotic and abiotic drivers on arthropod co-occurrence network topology in native forest remnants in the Azores"

### Supplementary Material 3

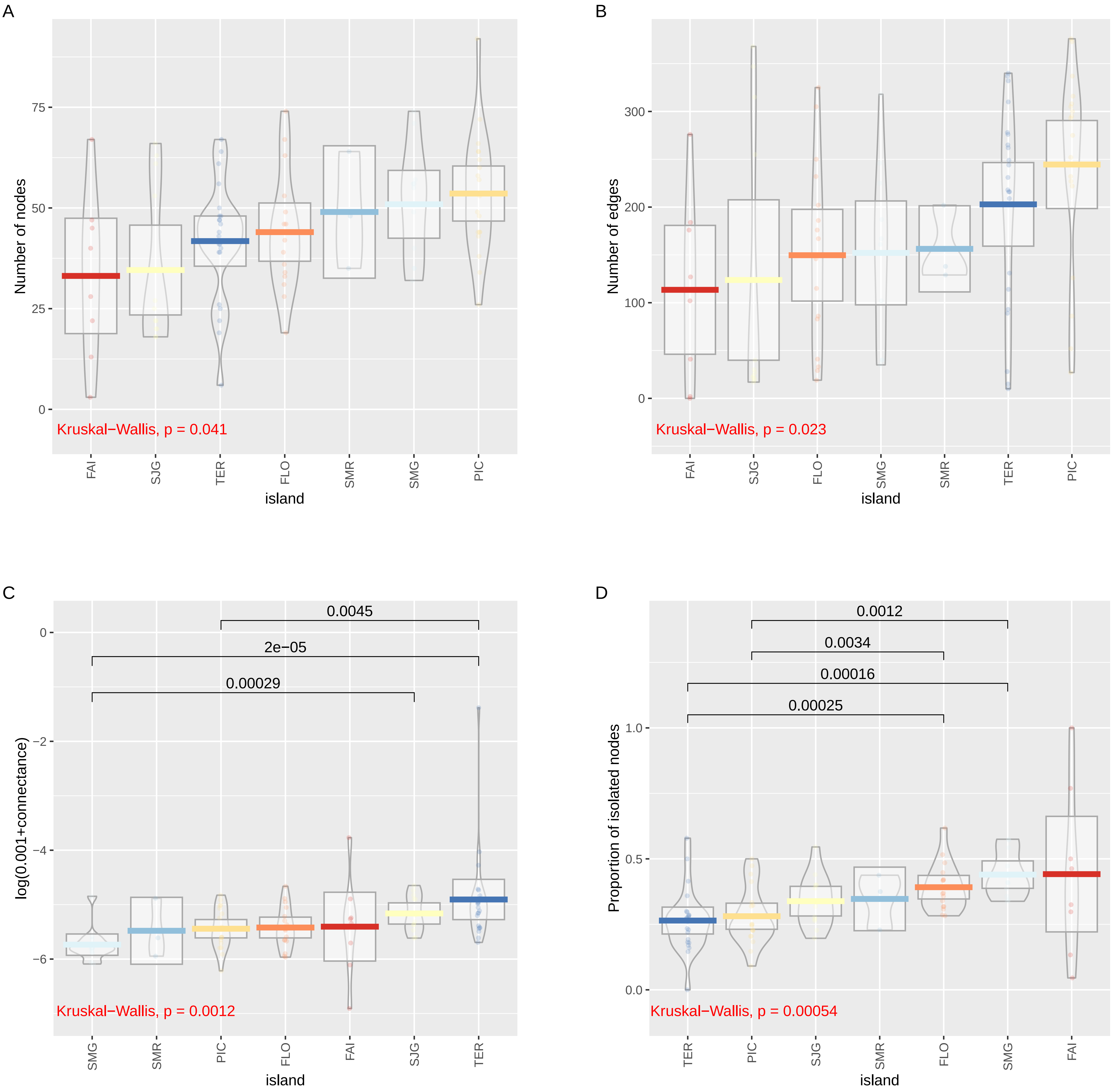

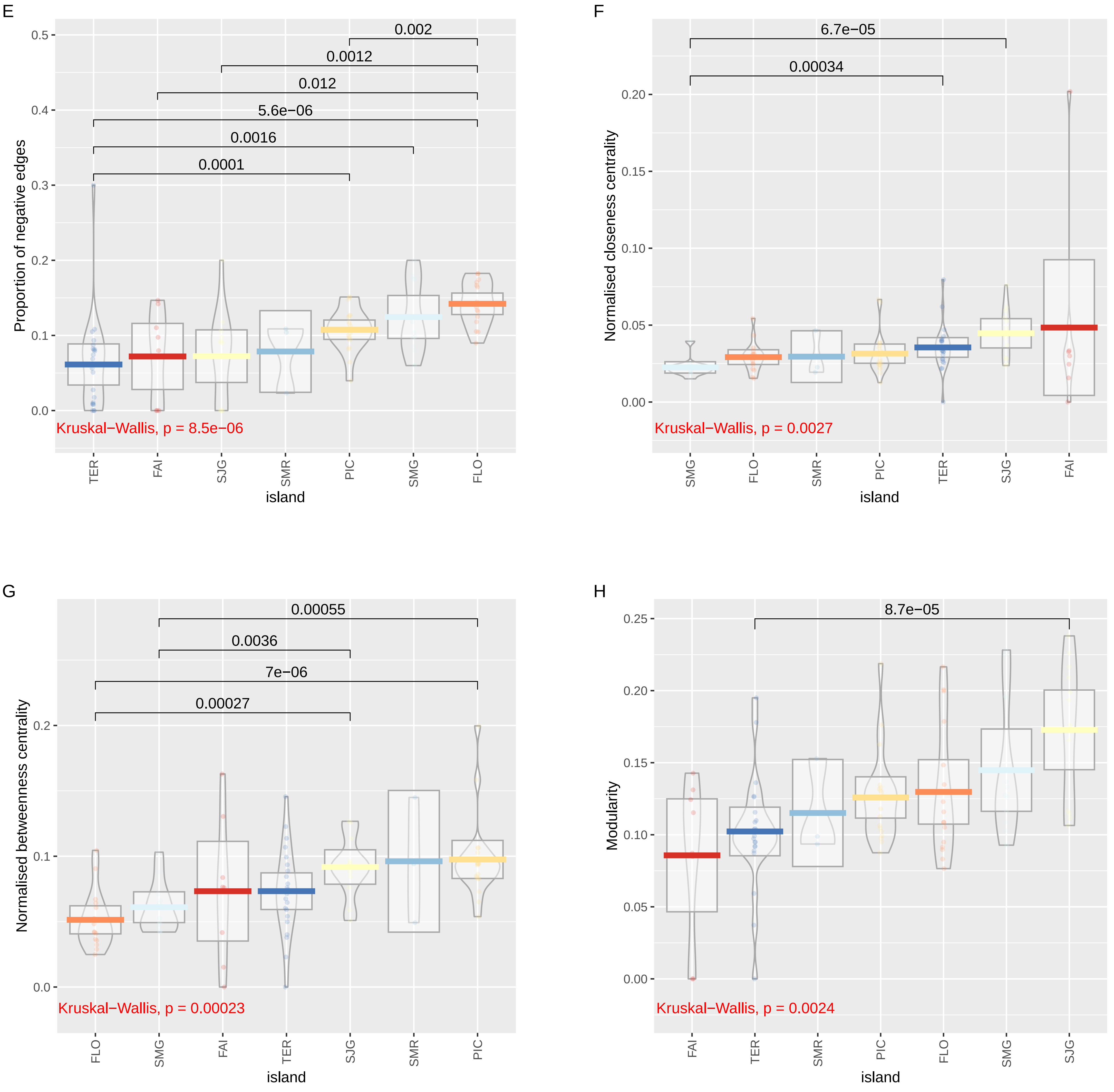

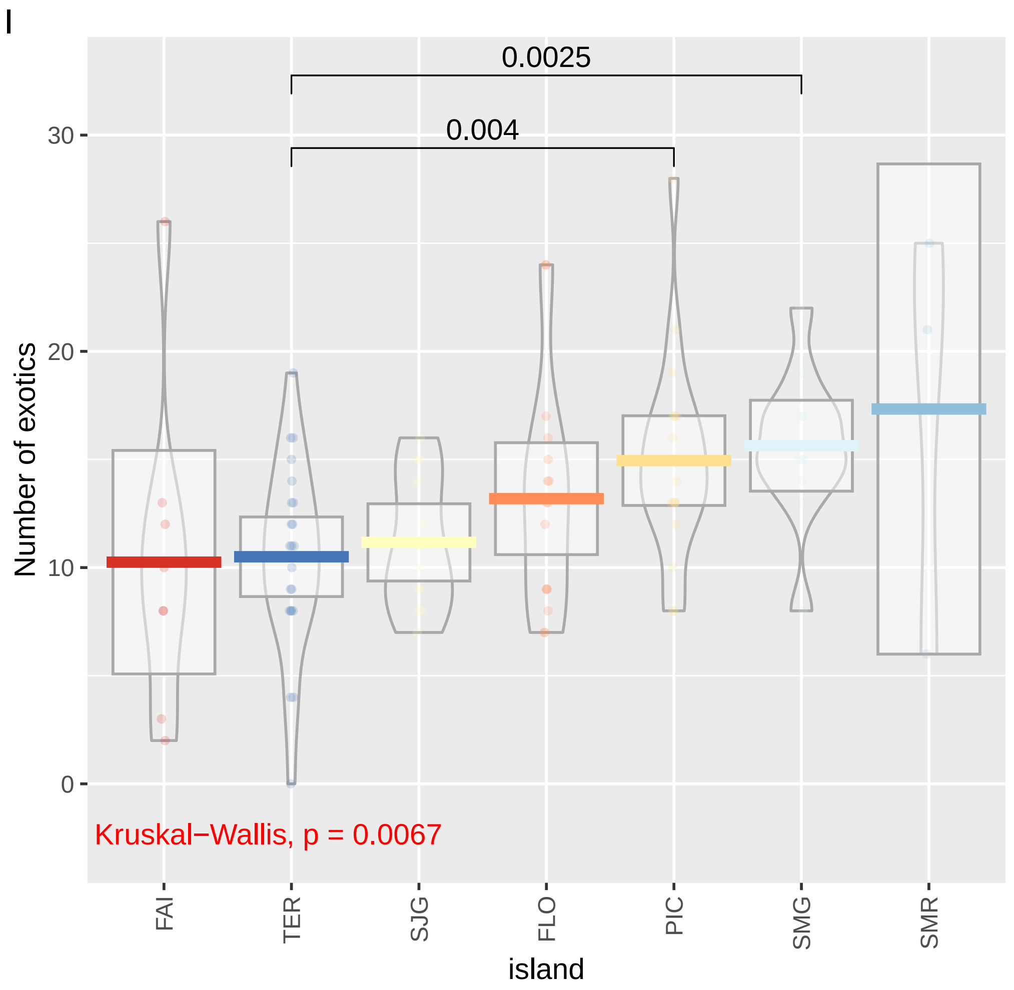

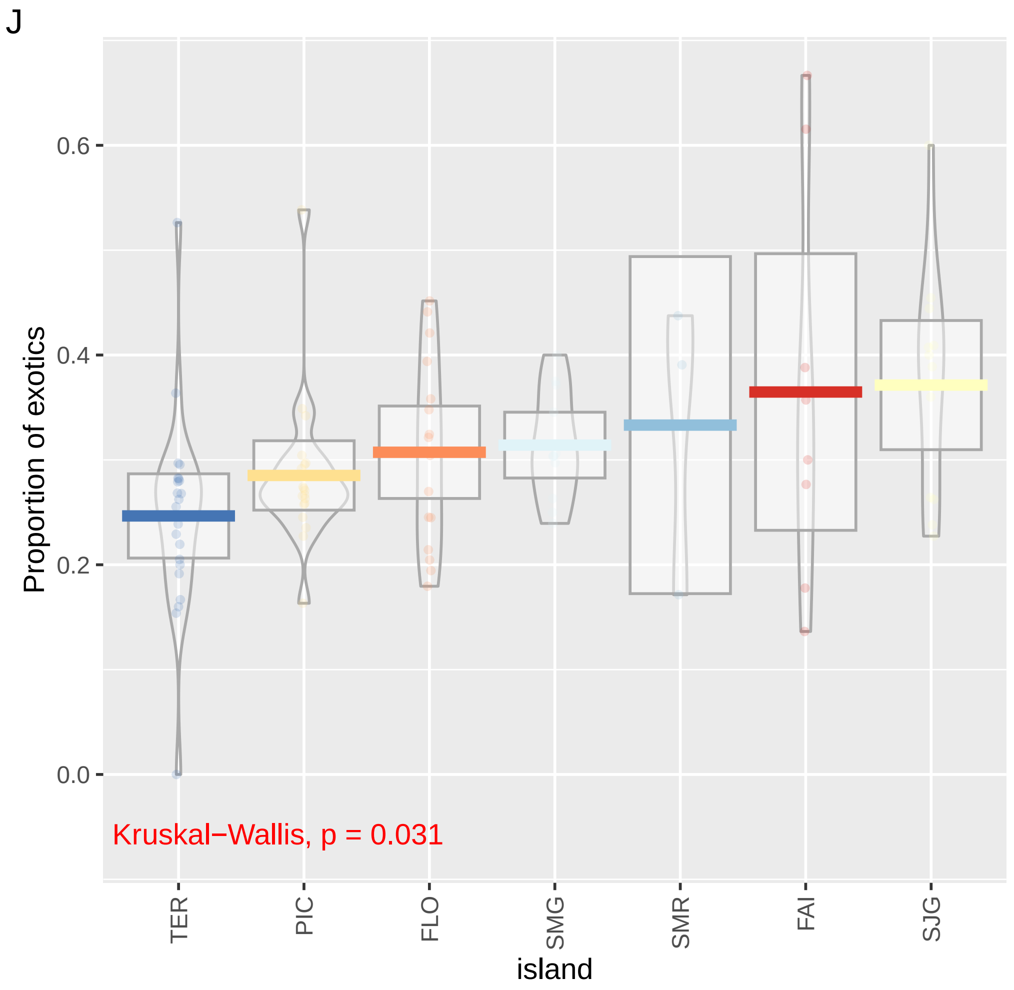

### Supplementary Material 5

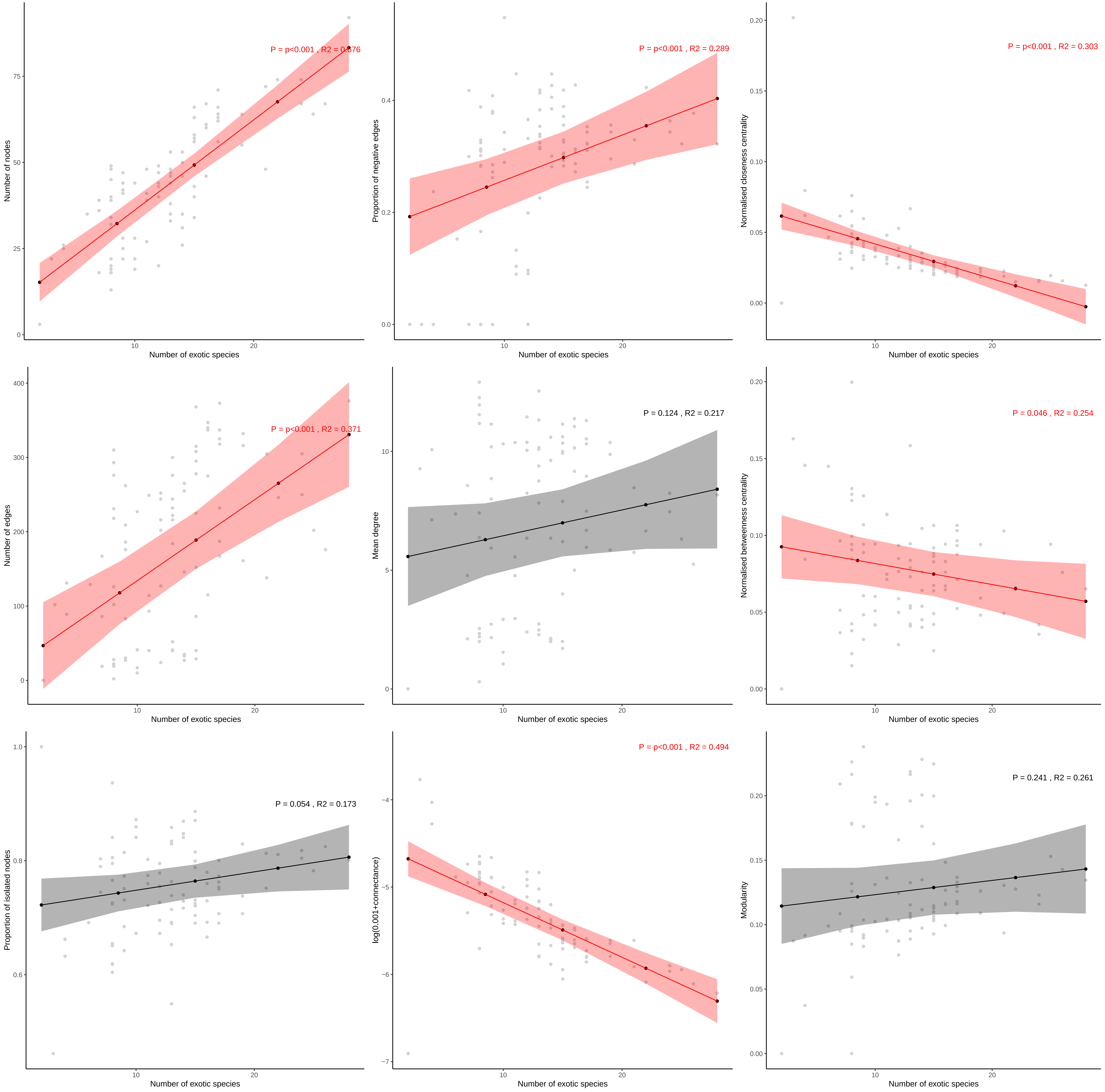

### Supplementary Material 6

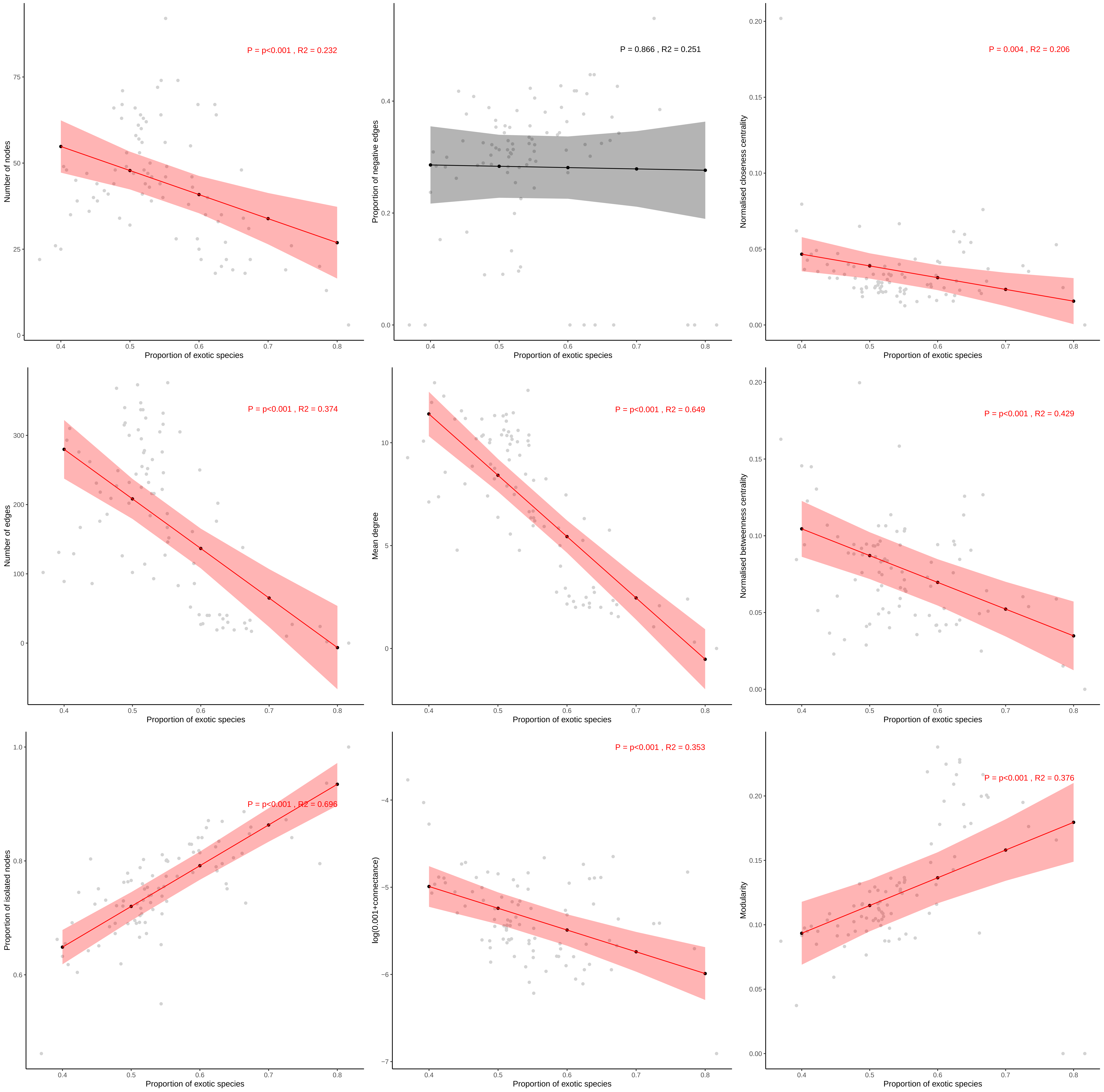

### Supplementary Material 7

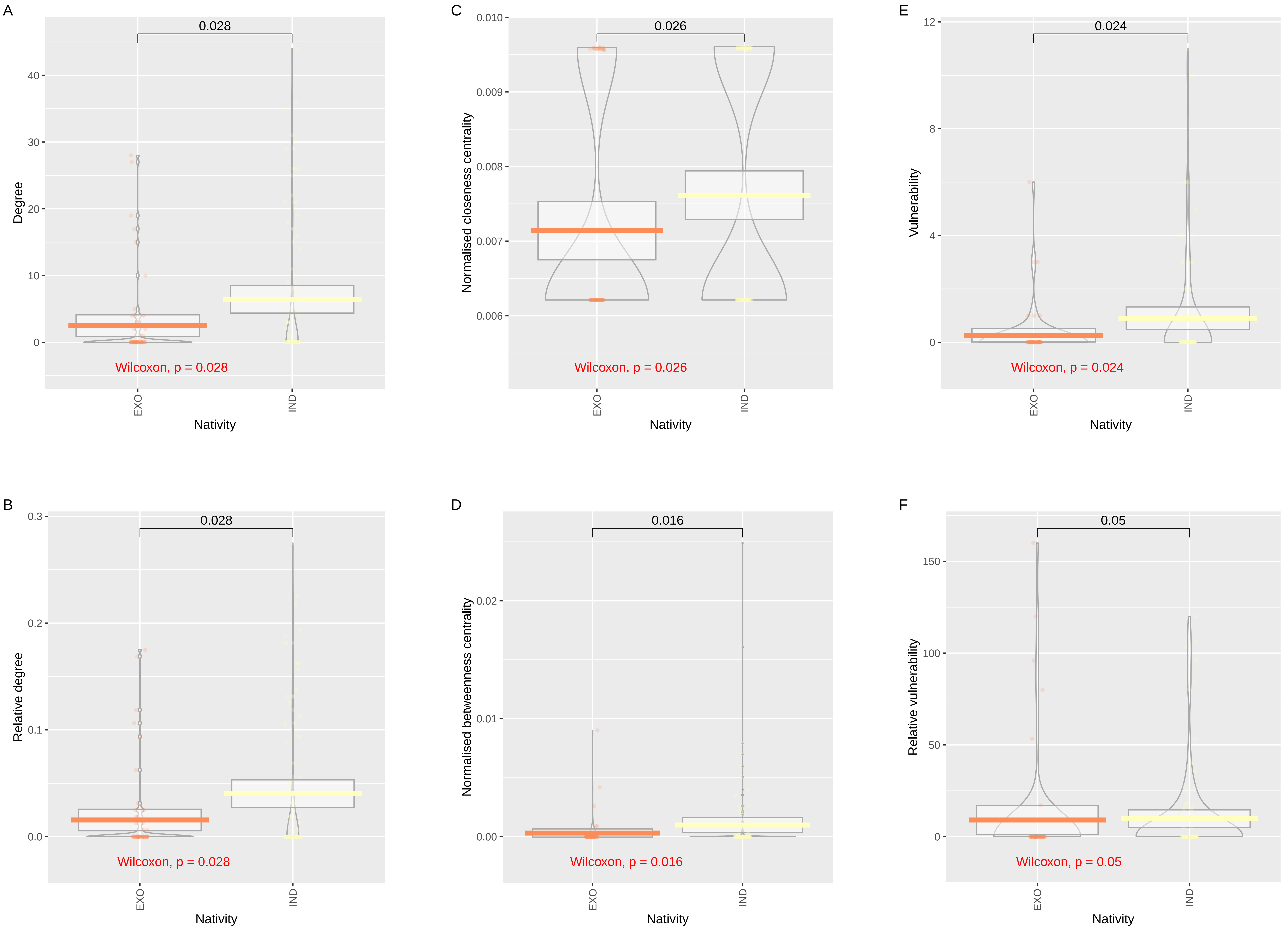

### Supplementary Material 8

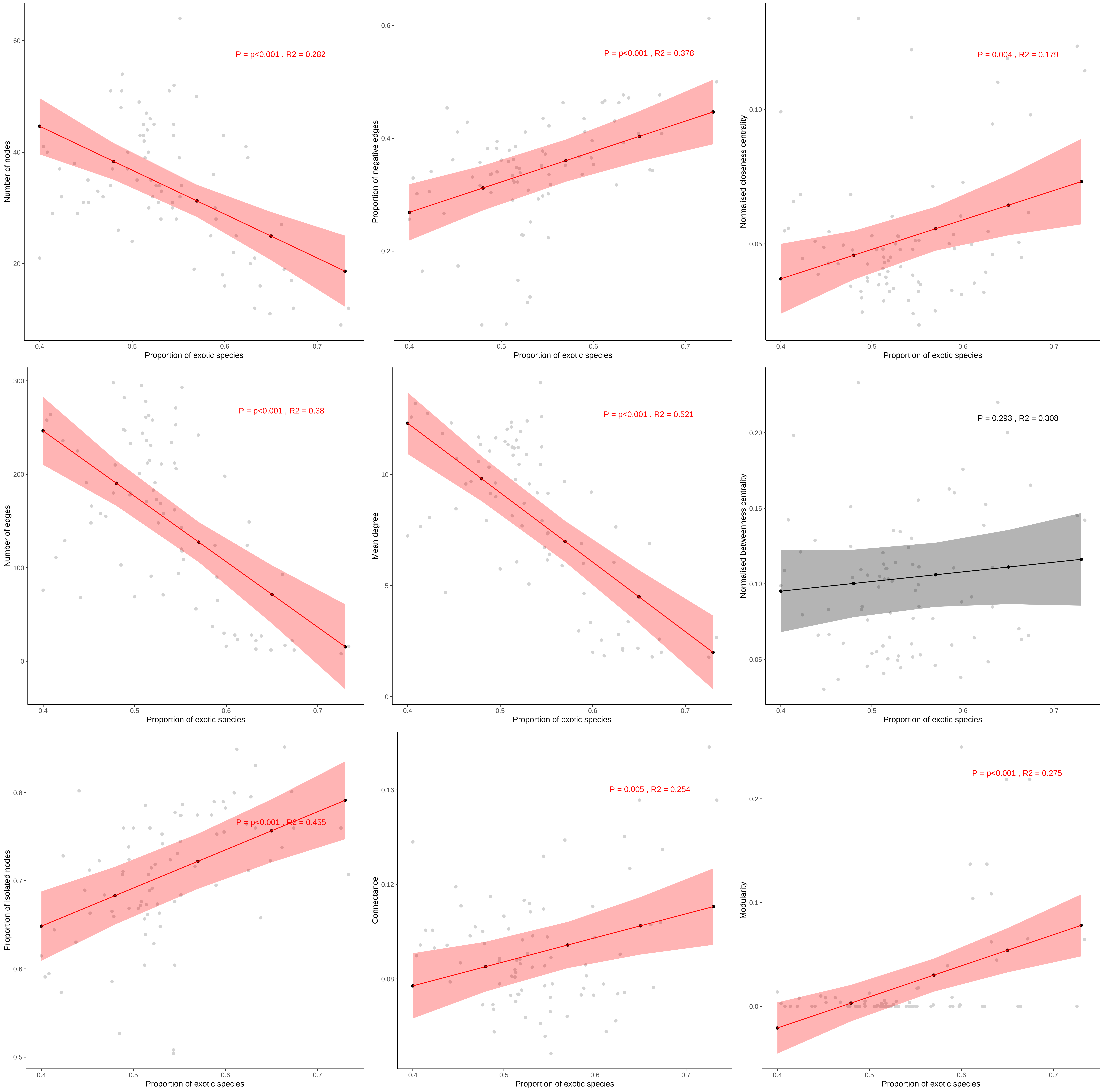

### Supplementary Material 9

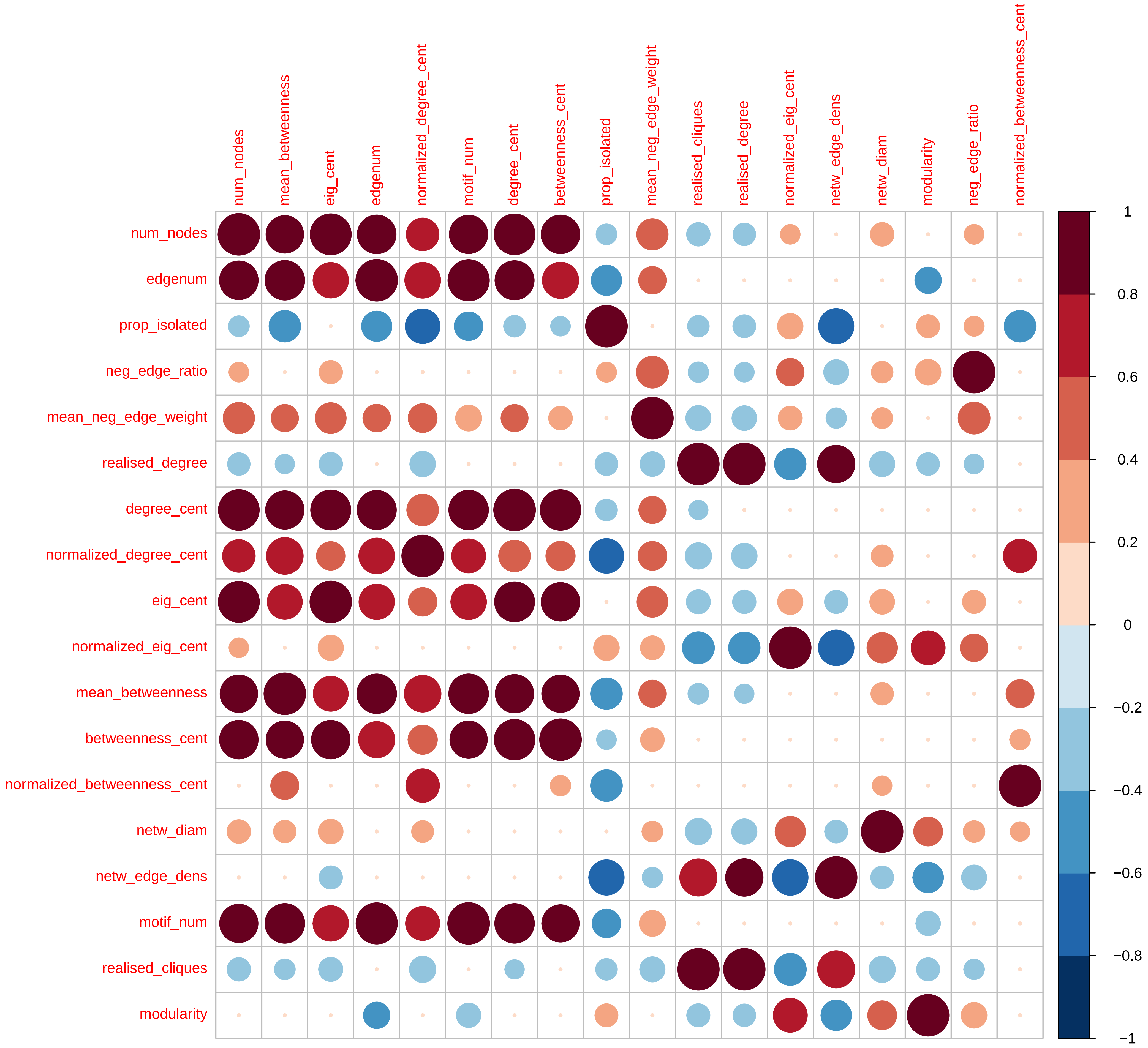

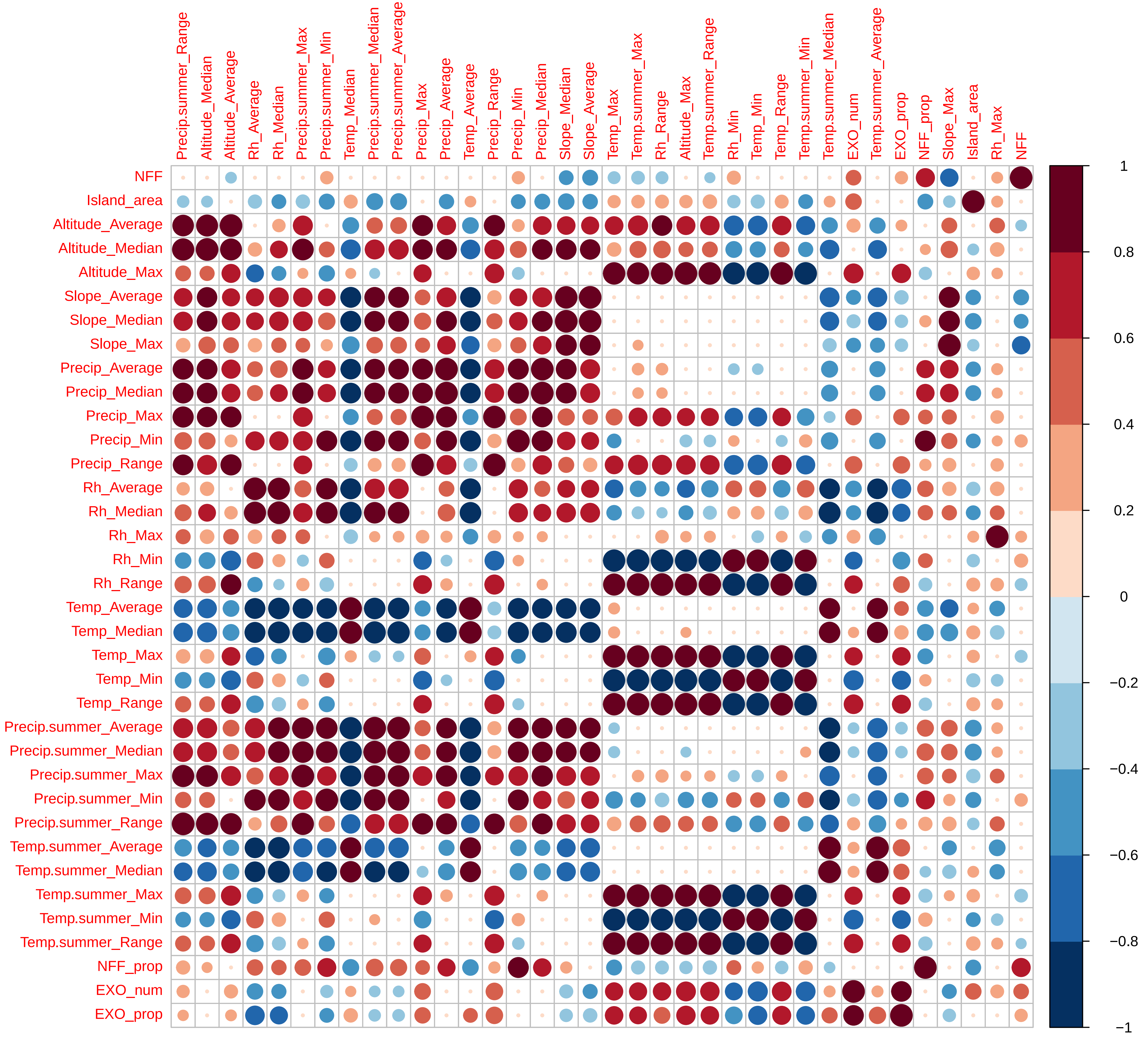

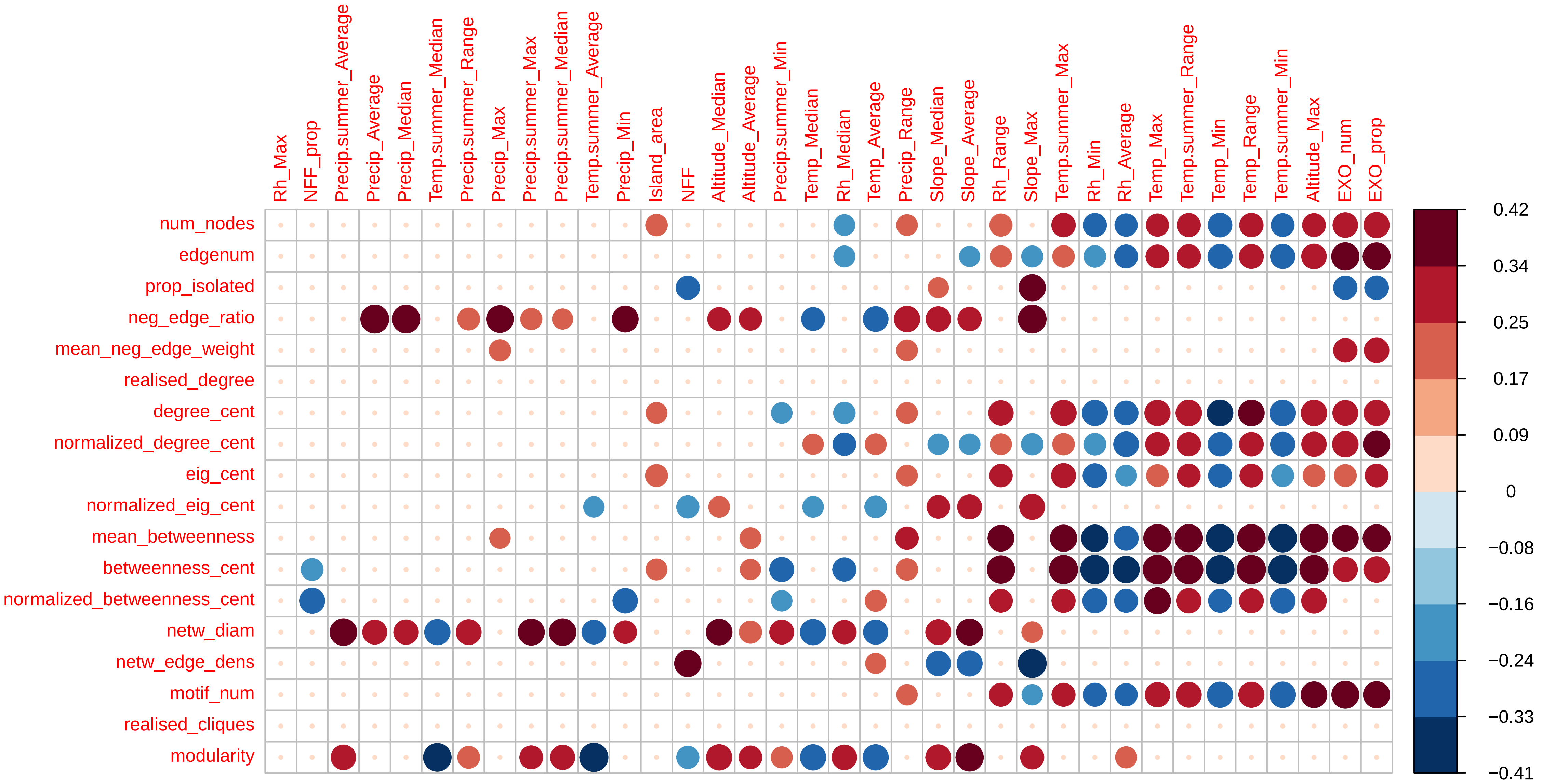
