## Supplementary Material 4 for "The influence of biotic and abiotic drivers on arthropod co-occurrence network topology in native forest remnants in the Azores"

**A) P-values of one-sided t-tests**

| Group | Test | E–E | E–I | E–N | I–I | I–N | N–N |
|---|---|---|---|---|---|---|---|
| **All links** | **Less** | <0.001 | 1 | <0.001 | 1 | 1 | <0.001 |
| | **Greater** | 1 | <0.001 | 1 | <0.001 | <0.001 | 1 |
| | **Two-sided** | <0.001 | <0.001 | <0.001 | <0.001 | <0.001 | <0.001 |
| **Negative links** | **Less** | 1 | 1 | <0.001 | 1 | 1 | <0.001 |
| | **Greater** | <0.001 | <0.001 | 1 | <0.001 | <0.001 | 1 |
| | **Two-sided** | <0.001 | <0.001 | <0.001 | <0.001 | <0.001 | <0.001 |

**B) Proportion test between exotic – native and native – native**

2-sample test for equality of proportions with continuity correction

X-squared = 9.457, df = 1, p-value = 0.001

Alternative hypothesis: less

95 percent confidence interval:

-1.000 -0.034

Sample estimates:

Prop 1 – Prop 2

0.090 0.166

**C) Proportion test between exotic – endemic and endemic – endemic**

2-sample test for equality of proportions with continuity correction

X-squared = 16.792, df = 1, p-value > 0.001

Alternative hypothesis: less

95 percent confidence interval:

-1.000 -0.065

Sample estimates:

Prop 1 – Prop 2

0.111 0.221

**D) Proportion test between exotic – indigenous and indigenous – indigenous**

**I-E + I-N and N+N + N-E + E-E**

2-sample test for equality of proportions with continuity correction

X-squared = 272.84, df = 1, p-value < 0.001

Alternative hypothesis: less

95 percent confidence interval:

-1.000 -0.538

Sample estimates:

Prop 1 – Prop 2

0.201 0.789
